# AI-Guided Design of Cyclic Peptide Binders Targeting TREM2 Using CycleRFdiffusion and Experimental Validation

**DOI:** 10.1101/2025.09.18.676322

**Authors:** Sungwoo Cho, Renjie Zhu, Katarzyna Kuncewicz, Hongliang Duan, Moustafa Gabr

## Abstract

Triggering receptor expressed on myeloid cells 2 (TREM2) plays a central role in regulating microglial function in the central nervous system and has emerged as a promising therapeutic target for Alzheimer’s disease. Despite advances in antibody-based therapeutics, small molecules and peptides capable of modulating TREM2 remain limited. Here, we present a cyclic peptide design pipeline that integrates CycleRFdiffusion, ProteinMPNN for sequence design, and HighFold for structural prediction and screening. Using the TREM2 structure as input, we generated and screened 1,500 peptide–target complexes, prioritizing four candidates that met structural and energetic criteria. Subsequent biophysical evaluation identified **TP4** as a TREM2 binder, demonstrating consistent binding in spectral shift, microscale thermophoresis, and surface plasmon resonance. Pharmacokinetic profiling indicated that **TP4** possesses favorable plasma stability and moderate metabolic stability, supporting its tractability for further optimization. This study establishes a generalizable framework for AI-driven cyclic peptide discovery and provides the first proof-of-concept demonstration of TREM2-targeted cyclic peptide binders.

---

Peptide-based therapeutics have gained significant attention as an intermediate modality between small molecules and biologics.^1^ Within this class, cyclic peptides have emerged as particularly attractive due to their constrained backbone conformations, which confer enhanced proteolytic stability, improved pharmacokinetic properties, and increased binding affinity to protein surfaces.^2^ Unlike linear peptides, cyclic peptides can effectively engage shallow or extended binding sites often found in protein–protein interactions (PPIs), making them compelling candidates for “undruggable” targets.^3^ Over the past two decades, several cyclic peptide drugs have reached clinical approval, demonstrating their potential as a therapeutic class for diverse indications including cancer, infectious disease, and autoimmune disorders.^3^ Despite these advances, rational discovery of cyclic peptides remains challenging, as the conformational space is vast and the prediction of stable binding poses requires methods that account for both sequence and topology.

TREM2 (triggering receptor expressed on myeloid cells 2) is a transmembrane receptor highly expressed on microglia, the resident immune cells of the central nervous system (CNS).^4,5^ Genetic studies have established strong associations between TREM2 variants and neurodegenerative diseases, particularly Alzheimer’s disease (AD), where loss-of-function mutations impair microglial activation, phagocytosis, and amyloid clearance.^6,7^ Consequently, pharmacological modulation of TREM2 is considered a promising therapeutic strategy to restore microglial function and potentially slow disease progression. To date, most drug discovery efforts targeting TREM2 have centered on monoclonal antibodies, several of which are advancing in clinical development.^8^ However, antibodies face limitations for CNS disorders, including poor blood–brain barrier (BBB) penetration, high production costs, and limited modes of administration.^9^ These drawbacks underscore the need for alternative modalities capable of targeting TREM2 with greater flexibility and accessibility.

The advent of artificial intelligence (AI)–driven protein design has provided powerful tools to explore chemical and structural spaces previously inaccessible through conventional medicinal chemistry. In this context, the emergence of RFdiffusion^10^ has marked a remarkable transition in protein design. Renowned for its high accuracy and reliability, this method has found extensive applications across diverse fields, including cancer immunotherapy, protease research, and the development of snake venom neutralizing proteins. Despite its remarkable achievements in binder design, the RFdiffusion model is currently suitable for linear peptides rather than cyclic formats. However, cyclic peptides, characterized by their unique cyclic backbone structures, possess distinct advantages over their linear counterparts^11-13^ Therefore, cyclic peptides represent promising candidates for therapeutic development, particularly in the context of challenging molecular interfaces.^14-16^ To address the unique challenges of cyclic peptide design, we engineered the RFdiffusion framework by modifying its input features specifically for cyclic peptide backbone generation, resulting in the CycleRFdiffusion model.^17^ Subsequently, we established a cyclic peptide binder design pipeline integrating CycleRFdiffusion (for backbone generation), ProteinMPNN (for sequence design), and HighFold^18,19^ (for cyclic peptide structure prediction). Finally, we applied an established scoring system to evaluate and prioritize the most promising cyclic peptide candidates. This integrative approach establishes a robust and comprehensive platform for the computational design and structural analysis of cyclic peptide binders.

To validate this framework, we applied it to the design of cyclic peptide binders targeting TREM2. Sequence design and structural prediction yielded a large ensemble of peptide–TREM2 complexes, which were subsequently filtered based on stringent structural confidence and interface energy metrics (Figure 1). Four high-quality peptide candidates were prioritized for experimental synthesis and evaluation. Finally, we performed biophysical validation using spectral shift, microscale thermophoresis (MST), and surface plasmon resonance (SPR) assays. Among the synthesized peptides, **TP4** emerged as a validated TREM2 binder, displaying consistent binding signals and yielding an equilibrium dissociation constant (Kd) in the sub-millimolar range. Although modest in affinity, this result provides the first demonstration of an AI-designed cyclic peptide capable of binding TREM2. Taken together, our study introduces a generalizable computational pipeline for cyclic peptide discovery and establishes proof-of-concept that small, AI-generated peptides can modulate CNS immune checkpoints, offering a potential alternative to antibody-based approaches.

**Figure 1.**
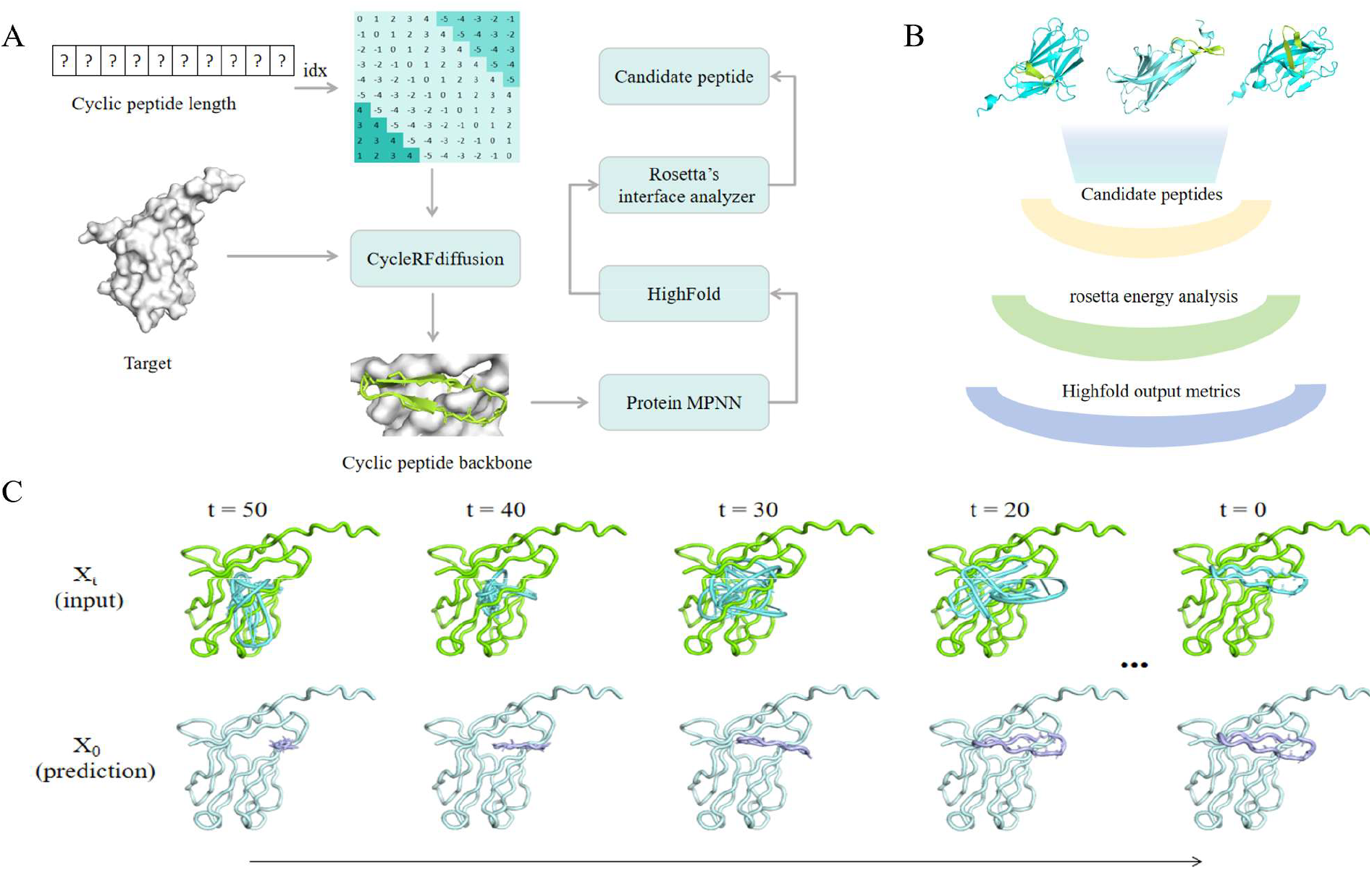
Workflow of Cyclic Peptide Binder Design and Screening. (**A**) CycleRFdiffusion generates cyclic peptide backbones for specific targets. These cyclic peptide backbones are then input into ProteinMPNN for sequence design. Subsequently, the designed sequences undergo structural prediction using HighFold. The final experimental candidates are obtained through screening using HighFold-based metrics and Rosetta’s Interface Analyzer.(**B**)Screening Process. The generated complexes are subjected to preliminary screening (screening criteria: iptm > 0.8, ipae < 0.3, plddt > 80,RMSD to input backbone <2 Å), followed by further screening using Rosetta’s Interface Analyzer (screening criterion: dGseparated/dSASA × 100 < 0). (**C**)A simple diagram of the denoising process (where X_t_ represents the noise input at the T-th time step, and X_0_ represents the denoised result predicted for the final time step at the T-th time step).

The cyclic peptide binder design targeting TREM2 was based on the experimentally observed structure (PDB: 6Y6C). Given that CycleRFdiffusion offers the option to incorporate hotspot residues in the design process, we implemented this design approach by using two different ways (Figure 1): (1) a hotspot-incorporated design and (2) a baseline design without hotspot residues. For the former, amino acid residues on the target chain in 6Y6C that were within 5 Å of the ligand chain were selected as hotspot residues (A59, A60, and A61). Subsequently, water molecules were removed from the 6Y6C structure, and only chain A was retained. This modified structure was then saved as a new PDB file, which served as the TREM2 target input for the CycleRFdiffusion model. This approach guaranteed the complete exclusion of any structural information from the original cyclic peptide ligand in the model input, facilitating plausible de novo generation of novel cyclic peptide backbones (Figure 1).

The TREM2 target was introduced into the CycleRFdiffusion model. The parameters for the generation of cyclic peptide backbones were configured as follows: backbone length was set to range from 6-16 residues, the number of diffusion steps was fixed at 50, and the quantity of generated backbones was specified as 30. Considering the two design types (with and without specified hotspot residues), the model ultimately yielded a total of 60 backbone design results. The output of CycleRFdiffusion consisted of cyclic peptide backbones that all residues being glycine (G).

ProteinMPNN was employed to design amino acid sequences based on three-dimensional protein structural features, including backbone atoms (N, Cα, C, O) and virtual Cβ positions. The CycleRFdiffusion-generated PDB files were processed in batch mode and used as input for ProteinMPNN, with the sampling temperature parameter fixed at 0.1 to ensure high-quality sequence predictions. For each backbone conformation, five distinct sequences were generated, resulting in a set of 300 designed sequences for subsequent evaluation.

The generated cyclic peptide sequences were then fed into HighFold to further assess the rationality of the designed backbones and sequences. HighFold incorporates a Cyclic Position Offset Encoding Matrix (CycPOEM) that is customized to the topological characteristics of cyclic peptides. This innovation ensures precise prediction of cyclic peptide structures and their interactions with target proteins. For each sequence, five predicted three-dimensional structures of the complexes were generated, leading a total of 1500 complex structures (Figure 2A-D).

**Figure 2.**
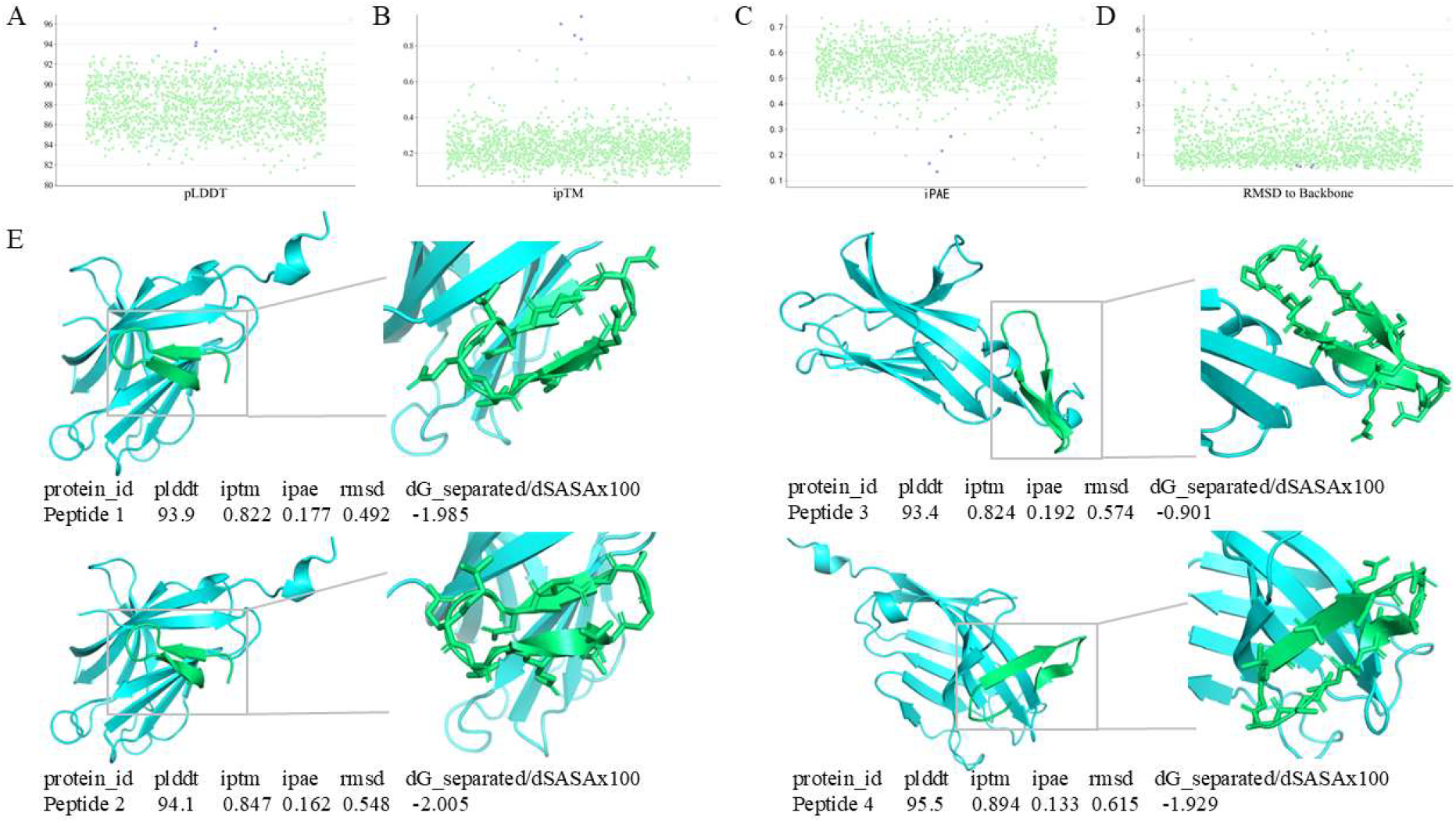
Design Metrics of Cyclic Peptides (Purple Dots Represent Filtered Cyclic Peptides) and Filtered Complex Structures. **(A)** pLDDT distribution of designed cyclic peptides. **(B)** ipTM distribution of designed cyclic peptides. **(C)** iPAE distribution of designed cyclic peptides. **(D)** RMSD to backbone distribution of designed cyclic peptides. **(E)** Detailed information about the cyclic peptide candidates (**TP1-TP4**).

The 1500 cyclic peptide-target complex structures were filtered using metrics derived from HighFold. This filtering step aimed to identify candidates for subsequent energy function calculations. These metrics have been validated in previous research and are recognized as reliable indicators for filtering in the realm of protein design.^20-22^ The selection criteria were established based on a previous design study,^23^ with design results featuring iPAE < 0.3 Å, ipTM > 0.8, RMSD to input backbone <2 Å, and pLDDT > 80 being selected. This process ultimately yielded four designs with high potential, all of which exhibited pLDDT values exceeding 93. Subsequently, Rosetta’s interface analysis tools were employed to perform energy minimization on these four designs and compute their respective energy functions. All four structures met the specified requirement (dGseparated/dSASA × 100 < 0), with three of them demonstrating dGseparated/dSASA × 100 < - 1.5. Consequently, these four sequences (Table 1) were selected for experiment validation (Figure 2E).

**Table 1.**
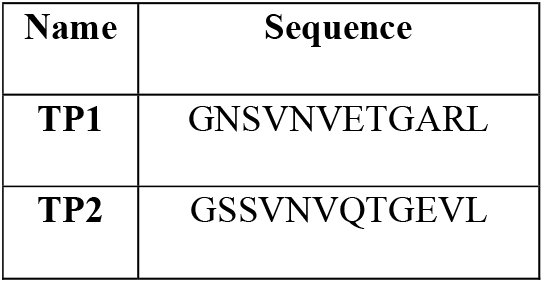
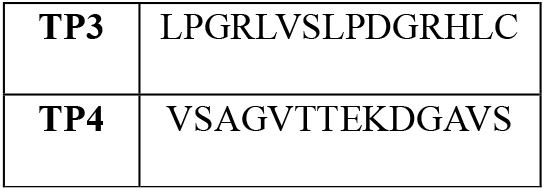
Sequences of the designed cyclic peptides (**TP1-TP4**) for further evaluation for TREM2 binding.

| Name | Sequence |
| --- | --- |
| TP1 | GNSVNVETGARL |
| TP2 | GSSVNVQTGEVL |
| TP3 | LPGRLVSLPDGRHLC |
| TP4 | VSAGVTTEKDGAVS |

The four cyclic peptide candidates (**TP1–TP4**) identified from computational screening were synthesized and evaluated using biophysical assays to assess their ability to bind TREM2. As shown in Figure 3, initial characterization by spectral shift analysis revealed measurable responses for **TP1, TP2**, and **TP4**, suggesting potential interactions with the TREM2 extracellular domain. However, **TP3** did not exhibit detectable binding under the tested conditions, consistent with its weaker computational scores. Based on the dose-response curves shown in Figure 3, the estimated Kd values for TREM2 binding affinity for peptides **TP1, TP2**, and **TP4** were 2.47 ± 1.21 mM, 1.54 ± 0.75, and 0.78 ± 0.13, respectively (Figure 3)\. These values indicate that **TP2** and **TP4** exhibit the most promising binding profiles, aligning with their strong computational predictions. Although the affinities remain in the millimolar range, they provide valuable starting scaffolds for further optimization. Importantly, these results establish proof-of-concept that AI-guided cyclic peptide design can yield experimentally validated binders against TREM2, a challenging neuroimmune target.

**Figure 3.**
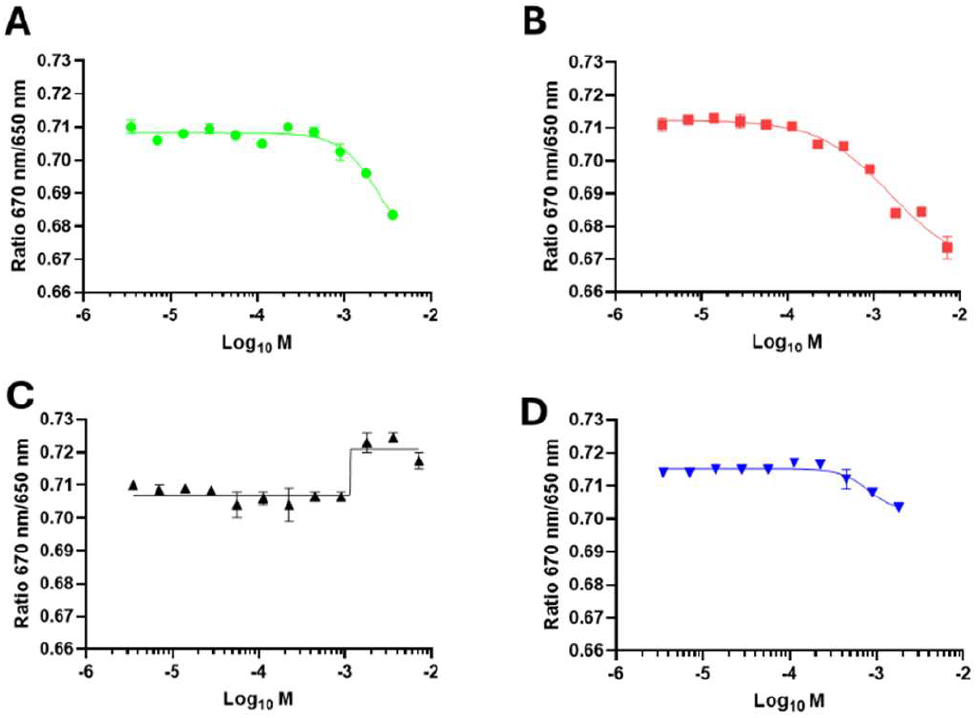
Biophysical validation of the four peptides using spectral shift analysis. Dose-response curves for **TP1 (A), TP2 (B), TP3 (C)**, and **TP4 (D)** in the spectral shift assays. Data represent mean ± SEM (n = 3).

To confirm these observations, the four peptides (**TP1-TP4**) were examined for TREM2 binding using MST. As shown in Figure 4, three peptides **TP1, TP2**, and **TP4** revealed dose-dependent changes in the Fnorm signal, which represents the normalized fluorescence intensity measured before and after thermophoretic movement of molecules in a microscopic temperature gradient. This normalization corrects for variations in initial fluorescence and allows direct comparison of bound versus unbound states. The concordance between MST and spectral shift results further strengthens the reliability of these peptides as TREM2 binders. In contrast, **TP3** did not yield a measurable MST response (Figure 4C), further supporting its weaker binding potential under these assay conditions.

**Figure 4.**
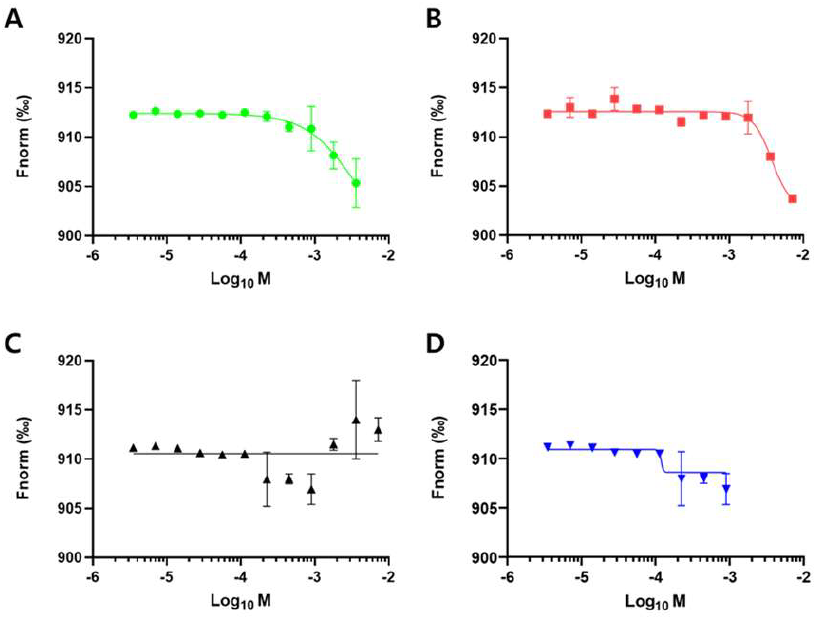
Dose-response curves for **TP1 (A), TP2 (B), TP3 (C)**, and **TP4 (D)** in the MST assays. Data represent mean ± SEM (n = 3).

Further investigation of **TP4** was performed using surface plasmon resonance (SPR) to validate its TREM2 binding affinity. SPR is a label-free technique that enables real-time monitoring of biomolecular interactions and provides kinetic information on binding events. SPR analysis demonstrated that **TP4** bound to TREM2 in a dose-dependent manner, with increasing concentrations producing progressively higher response signals (Figure 5). Although precise affinity constants could not be determined with confidence, the observed sensorgrams confirmed that **TP4** is capable of engaging TREM2 under real-time kinetic conditions. These findings indicate that, **TP4** retains measurable binding activity toward TREM2 as detected by spectral shift, MST, and SPR.

**Figure 5.**
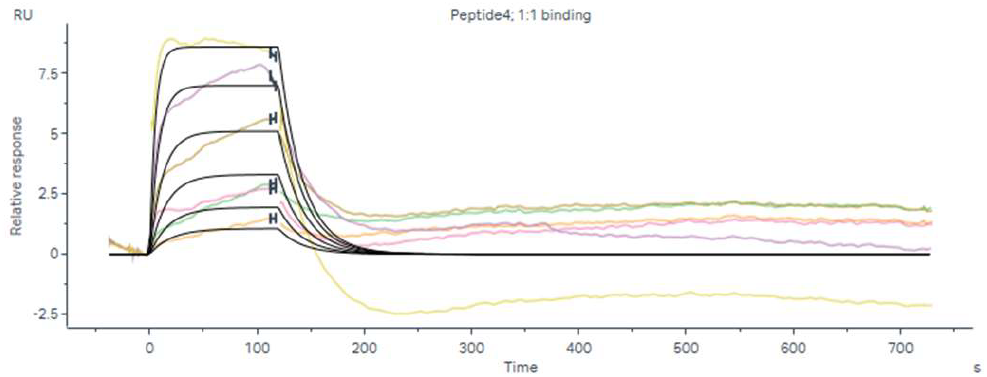
Sensorgram of **TP4** interacting with the TREM2 protein using SPR. TREM2 was immobilized on SA chip. Running buffer: PBS-P. **TP4** concentration: 3.75 – 60 µM (2-fold dilution).

To gain preliminary insights into the drug-like properties of the designed cyclic peptides, we next evaluated the in vitro pharmacokinetic profile of **TP4**. Such analyses are critical for assessing the developability of peptide scaffolds, as stability in biological matrices, lipophilicity, permeability, and metabolic turnover strongly influence their potential as therapeutic agents. As summarized in Table 2, **TP4** displayed a short half-life under simulated gastric conditions 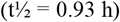 but was more stable in simulated intestinal fluid 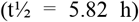, consistent with the protective effect of backbone cyclization at neutral pH. In human plasma, **TP4** retained 83.1% of its integrity after 1 h, highlighting good resistance to proteolysis. **TP4** exhibited a LogD_7.4_ of 1.03, indicative of a balanced hydrophilic–lipophilic profile, and demonstrated low-to-moderate permeability across Caco-2 cells ratio 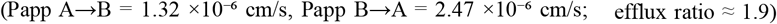, in line with expectations for a cyclic peptide. Finally, **TP4** showed moderate metabolic stability in rat liver microsomes 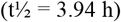. Collectively, these results suggest that **TP4** combines reasonable plasma and intestinal stability with modest permeability, supporting its potential as a tractable lead scaffold for further optimization

**Table 2.** *In vitro* pharmacokinetic profile of TP4.

| Test | TP4 |
| --- | --- |
| Stability in simulated fluids ($t_{1/2}$, h) | |
| Gastric (pH 1.6) | 0.93 |
| Intestinal (pH 6.5) | 5.82 |
| Stability in human plasma (% remaining at 1 h) | 83.1 |
| $\text{LogD}_{7.4}^a$ | 1.03 |
| CACO cell permeability<sup>b</sup> |  |
| $P_{app} A \rightarrow B$ ($10^{-6}$ cm/s) | 1.32 |
| $P_{app} B \rightarrow A$ ($10^{-6}$ cm/s) | 2.47 |
| Stability in rat liver microsomes ($t_{1/2}$, h) | 3.94 |

<sup>a</sup>Determined by miniaturized shake-flask procedure at pH7.4.
<sup>b</sup>Permeation through a Caco-2 cell monolayer was assessed in the absorptive apical to basolateral (a→b) and secretory basolateral to apical (b→a).

In summary, we established and experimentally validated an AI-guided pipeline for the discovery of cyclic peptide binders targeting TREM2. By integrating CycleRFdiffusion for backbone generation, ProteinMPNN for sequence design, and HighFold for structural assessment, we generated a diverse set of peptide– TREM2 complexes and prioritized candidates based on rigorous structural and energetic criteria. Four cyclic peptides were synthesized, among which **TP4** consistently demonstrated binding activity across spectral shift, MST, and SPR assays. Importantly, **TP4** also exhibited a favorable pharmacokinetic profile, with stability in intestinal fluid and plasma, moderate microsomal stability, and measurable though limited permeability. These results represent the first proof-of-concept demonstration of AI-designed cyclic peptides capable of binding TREM2 and provide a tractable starting point for optimization toward higher affinity and improved drug-like properties. More broadly, the study illustrates how AI-driven design can be leveraged to expand therapeutic modalities beyond antibodies and small molecules, enabling systematic exploration of cyclic peptides as potential immunomodulators for CNS targets such as TREM2.

## Data Availability

Data will be made available on request.

## References and notes

1. Sharma K, Sharma KK, Sharma A, Jain R. Peptide-based drug discovery: Current status and recent advances. Drug Discov Today. 2023, 28(2), 103464

2. Ji X, Nielsen AL, Heinis C. Cyclic Peptides for Drug Development. Angew Chem Int Ed Engl. 2024, 63(3), e202308251.

3. Ramadhani D, Maharani R, Gazzali AM, Muchtaridi M. Cyclic Peptides for the Treatment of Cancers: A Review. Molecules. 2022, 27(14), 4428.

4. Pocock J, Vasilopoulou F, Svensson E, Cosker K. Microglia and TREM2. Neuropharmacology. 2024, 257, 110020.

5. Yeh FL, Hansen DV, Sheng M. TREM2, Microglia, and Neurodegenerative Diseases. Trends Mol Med. 2017, 23(6), 512–533.

6. Qin Q, Teng Z, Liu C, Li Q, Yin Y, Tang Y. TREM2, microglia, and Alzheimer’s disease. Mech Ageing Dev. 2021, 195, 111438.

7. Hou J, Chen Y, Grajales-Reyes G, Colonna M. TREM2 dependent and independent functions of microglia in Alzheimer’s disease. Mol Neurodegener. 2022, 17(1), 84.

8. Schlepckow K, Morenas-Rodríguez E, Hong S, Haass C. Stimulation of TREM2 with agonistic antibodies-an emerging therapeutic option for Alzheimer’s disease. Lancet Neurol. 2023, 22(11), 1048–1060.

9. Razpotnik R, Novak N, Čurin Šerbec V, Rajcevic U., Targeting Malignant Brain Tumors with Antibodies. Front Immunol. 2017, 8, 1181.

10. Watson J L, Juergens D, Bennett N R, et al. De novo design of protein structure and function with RFdiffusion[J]. Nature, 2023, 620(7976): 1089–1100.

11. Buckton, L. K.,, Rahimi, M. N.,, McAlpine, S. R. Cyclic Peptides as Drugs for Intracellular Targets: The Next Frontier in Peptide Therapeutic Development. Chemistry 2021, 27 (5), 1487– 1513.

12. Haberman, V. A.,, Fleming, S. R.,, Leisner, T. M.,, Puhl, A. C.,, Feng, E.,, Xie, L.,, Chen, X.,, Goto, Y.,, Suga, H.,, Parise, L. V. Discovery and Development of Cyclic Peptide Inhibitors of CIB1. ACS Med. Chem. Lett. 2021, 12 (11), 1832– 1839.

13. Zhang, H.,, Chen, S. Cyclic peptide drugs approved in the last two decades (2001–2021). RSC Chem. Biol. 2022, 3 (1), 18– 31.

14. Marsault, E.,, Peterson, M. L. Practical Medicinal Chemistry with Macrocycles; Wiley Online Library, 2017.

15. Cardote, T. A. F.,, Ciulli, A. Cyclic and macrocyclic peptides as chemical tools to recognise protein surfaces and probe protein– protein interactions. ChemMedChem 2016, 11 (8), 787– 794.

16. Qian, Z.,, Dougherty, P. G.,, Pei, D. Targeting intracellular protein– protein interactions with cell-permeable cyclic peptides. Curr. Opin. Chem. Biol. 2017, 38, 80– 86.

17. Zhang C, Xu Z, Lin K, et al. CycleDesigner: Leveraging CycRFdiffusion and HighFold to Design Cyclic Peptide Binders for Specific Targets[J]. Journal of Chemical Information and Modeling, 2025, 65(12): 6155–6165.

18. Evans, R.,, O’Neill, M.,, Pritzel, A.,, Antropova, N.,, Senior, A.,, Green, T.,, Žídek, A.,, Bates, R.,, Blackwell, S.,, Yim, J. Protein complex prediction with AlphaFold-Multimer. biorxiv 2021, 2021.2010. 2004.463034.

19. Zhang, C.,, Zhang, C.,, Shang, T.,, Zhu, N.,, Wu, X.,, Duan, H. HighFold: accurately predicting structures of cyclic peptides and complexes with head-to-tail and disulfide bridge constraints. Brief Bioinform 2024, 25 (3), bbae215.

20. Chae, J.,, Wang, Z.,, Gul, I.,, Ji, J.,, Chen, Z.,, Qin, P. pLDDT-Predictor: High-speed Protein Screening Using Transformer and ESM2, 2024, 2410.21283. arXiv.org e-Print archive.

21. Guo, X.-Y.,, Li, Y.-F.,, Liu, Y.,, Pan, X.,, Shen, H.-B. ProtDAT: A Unified Framework for Protein Sequence Design from Any Protein Text Description, 2024, 2412.04069. arXiv.org e-Print archive.

22. Xu, W.,, Wu, Z.,, Zhang, C.,, Zhu, C.,, Duan, H. PepCARES: A Comprehensive Advanced Refinement and Evaluation System for Peptide Design and Affinity Screening. ACS Omega 2024, 9 (46), 46429– 46438.

23. Pacesa, M.,, Nickel, L.,, Schellhaas, C.,, Schmidt, J.,, Pyatova, E.,, Kissling, L.,, Barendse, P.,, Choudhury, J.,, Kapoor, S.,, Alcaraz-Serna, A. BindCraft: one-shot design of functional protein binders. bioRxiv 2024.

