## Supporting Information for "AI-Guided Design of Cyclic Peptide Binders Targeting TREM2 Using CycleRFdiffusion and Experimental Validation"

*Electronic Supplementary Information*

| **Contents** |  |
| --- | --- |
| Experimental  References | S2  S4 |
| MS and HPLC data for the synthesized peptides | S5 |

**Experimental:**

**1. Synthesis of the Peptides:**

The peptides were synthesized on 2-CL resin, using standard Fmoc synthesis protocol with DIC/Cl-HOBt coupling, on an APEX 396 automatic synthesizer. The resin was swollen in DMF for 30 min, treated with 20v% Piperidine-DMF for 8 minutes at 50°C to remove the Fmoc protecting group and washed with DMF for three times. For the coupling reaction, the resin was added with Fmoc-protected amino acid, Cl-HOBt, DIC and NMP. The mixture was vortexed for 20 minutes at 50°C. Afterwards, the resin was washed with DMF once. The cycle of deprotection and coupling steps was repeated until the last amino acid residue was assembled. After the final Fmoc protecting group was removed, the resin was treated with 20v% acetic Anhydride-NMP for 20 minutes. The resin was then washed with DMF, DCM and dried with air. The peptides were cleaved using a TFA cocktail (95v%TFA, 2.5v%water and 2.5v%TIS) for three hours. Crude peptides were precipitated by adding ice-chilled anhydrous ethyl ether, washed with anhydrous ethyl ether three times, and freeze-dried. After cyclization, the crude peptides were loaded onto a prep-HPLC column and purified with a gradient of 10%-55%B within 45 minutes at a flow rate of 12 ml/min. The peptides were analyzed by LC-MS and confirmed to have >95% HPLC purity, and freeze-dried.

**2. Spectral Shift Assay**

Spectral shift measurements were performed using a Dianthus NT.23PicoDuo instrument (NanoTemper Technologies, Munich, Germany). TREM2 protein was labeled with RED-tris-NTA 2nd Generation dye using the His-Tag Labeling Kit (NanoTemper Technologies) according to the manufacturer's protocol. All binding experiments were conducted in PBST buffer containing 154 mM NaCl, 5.6 mM Na₂HPO₄, 1.05 mM KH₂PO₄, pH 7.4, and 0.005% Tween-20. For binding measurements, labeled TREM2 protein was used at a final concentration of 10 nM. Test peptides were serially diluted in PBST buffer and incubated with labeled TREM2 for 10 minutes at room temperature (22-25°C) prior to analysis. The Dianthus instrument was configured with the following parameters: 85% LED excitation power, picomolar detector sensitivity disabled, and laser on-time of 5 seconds. Data analysis was performed using Dianthus Analysis software (NanoTemper Technologies) for initial processing, followed by curve fitting and statistical analysis using GraphPad Prism 10.0 (GraphPad Software, San Diego, CA, USA).

**3. Microscale Thermophoresis (MST) Binding Assay**

Binding affinity measurements were performed using microscale thermophoresis (MST) on a Monolith NT.115 system (NanoTemper Technologies, Munich, Germany). TREM2 was labeled using the RED-tris-NTA His-tag labeling kit (NanoTemper Technologies) according to the manufacturer's instructions. MST measurements were conducted using different buffer conditions optimized for each protein: TREM2 assays were performed in PBS buffer (pH 7.4) containing 0.005% Tween-20. Labeled protein was used at a final concentration of 40 nM and incubated with serially diluted test peptides for 10 minutes at room temperature (22-25°C) prior to measurement. MST experiments were performed using standard capillaries with the following instrument parameters: red filter set, 100% LED power, and medium MST power. Thermophoresis was monitored for 20 seconds with an additional 5-second delay. Data analysis was conducted using MO.Affinity Analysis software (NanoTemper Technologies) for initial processing, followed by curve fitting using GraphPad Prism 10.0 (GraphPad Software, San Diego, CA, USA).

**4. Surface plasmon resonance (SPR)**

The interaction between **TP4** and the human TREM2 protein was evaluated using surface plasmon resonance (SPR) on a Biacore 8K instrument (Cytiva). Biotinylated TREM2 (Cat. No. 11084-H49H-B, Sino Biological) was immobilized on an SA Sensor Chip (Cytiva) in PBS-P buffer (0.2 M phosphate buffer, 27 mM KCl, 1.37 M NaCl, and 0.5% Surfactant P20, pH 7.4; Cytiva). Before protein immobilization, the chip surface was conditioned with three 1-minute injections of 1 M NaCl in 50 mM NaOH. Following immobilization, the surface was washed sequentially with 50% (v/v) isopropanol in water, 1 M NaCl, and 50 mM NaOH to remove weakly or non-specifically bound material. The system was then equilibrated in running buffer until a stable baseline was established. Serial concentrations of **TP4** (3.75 – 60 µM, 2-fold dilution) were prepared in PBS-P buffer and injected over the immobilized TREM2 surface. Experiments were carried out at 25°C with a flow rate of 30 μL/min, a contact time of 120 s, and a dissociation phase of 600 s. After each run, the surface was washed with 50% DMSO. Sensorgram was generated after subtracting background response signal from a reference flow cell and from a control experiment with buffer injection. Interaction was investigated at least in triplicate. Data analysis was performed using Biacore™ Insight Evaluation Software (Cytiva). A 1:1 binding model provided the best fit to the experimental data in single-cycle kinetic analysis.

**5. Pharmacokinetic profiling**

The assays for determining pharmacokinetic profile were performed as previously described.^1^

• Stability in intestinal simulated fluids (t1/2, min) was evaluated using erythromycin as a reference compound, t1/2, min = 453 min.^1^

• Stability in rat liver microsomes was evaluated using verapamil as a reference compound, t1/2, min = 6.3 min.^1^

• Human plasma stability was evaluated using propantheline as a reference compound, t1/2, min = 2.6 min.^1^

**References:**

1. Abdel-Rahman SA, Ovchinnikov V, Gabr MT. Structure-Based Rational Design of Constrained Peptides as TIM-3 Inhibitors. *ACS Med Chem Lett*. 2024;15(6):806-813. doi:10.1021/acsmedchemlett.3c00567

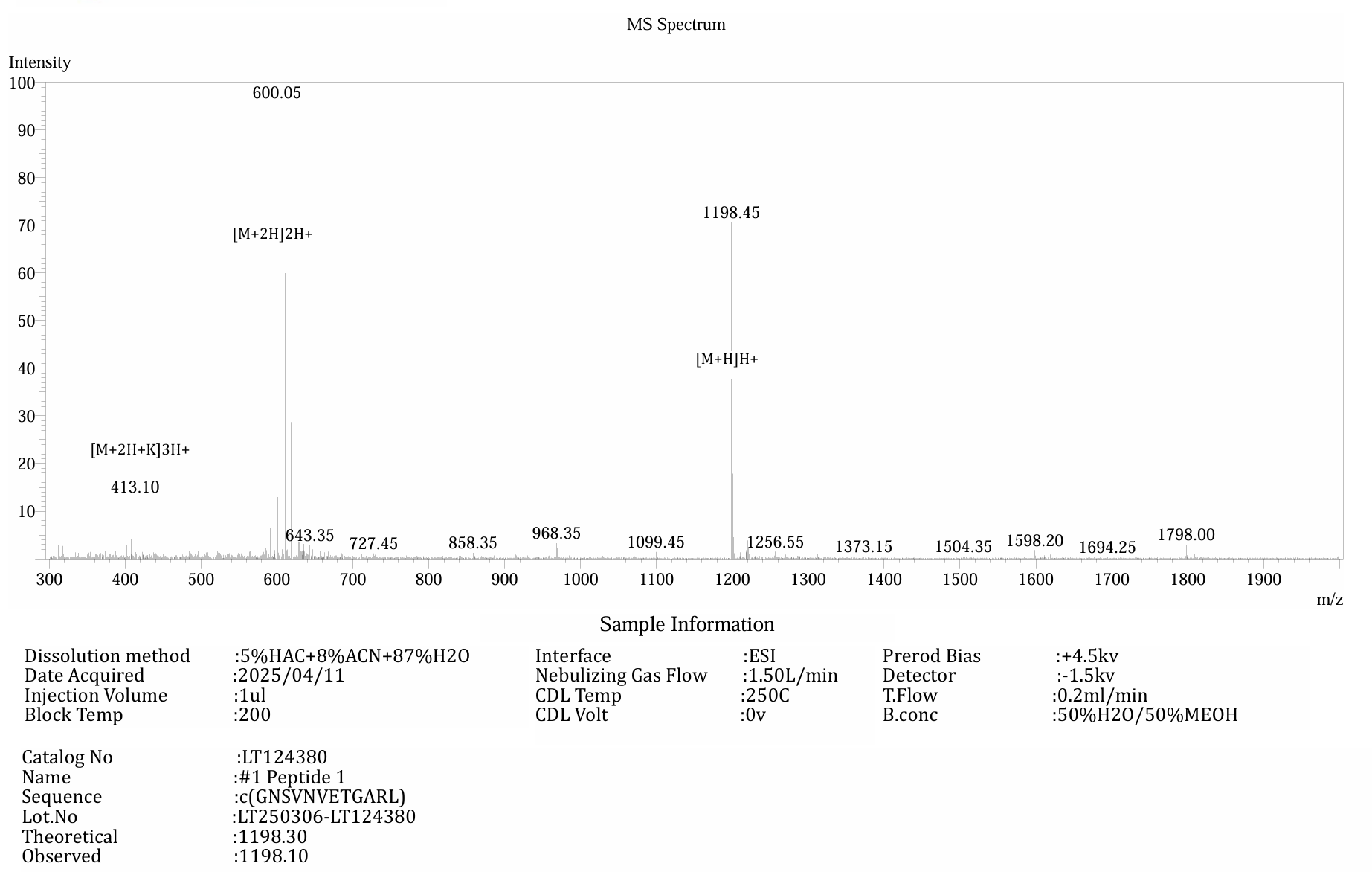

**Figure S1**. MS spectrum of **TP1**.

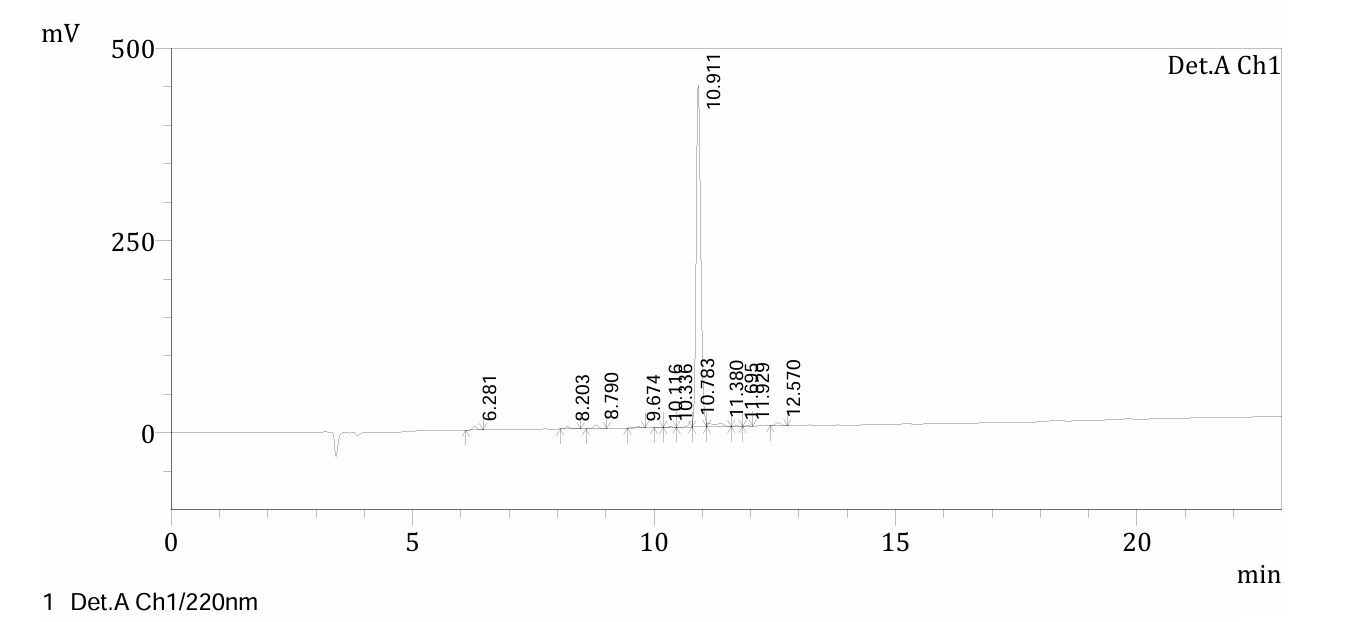

**Figure S2**. HPLC trace of **TP1**.

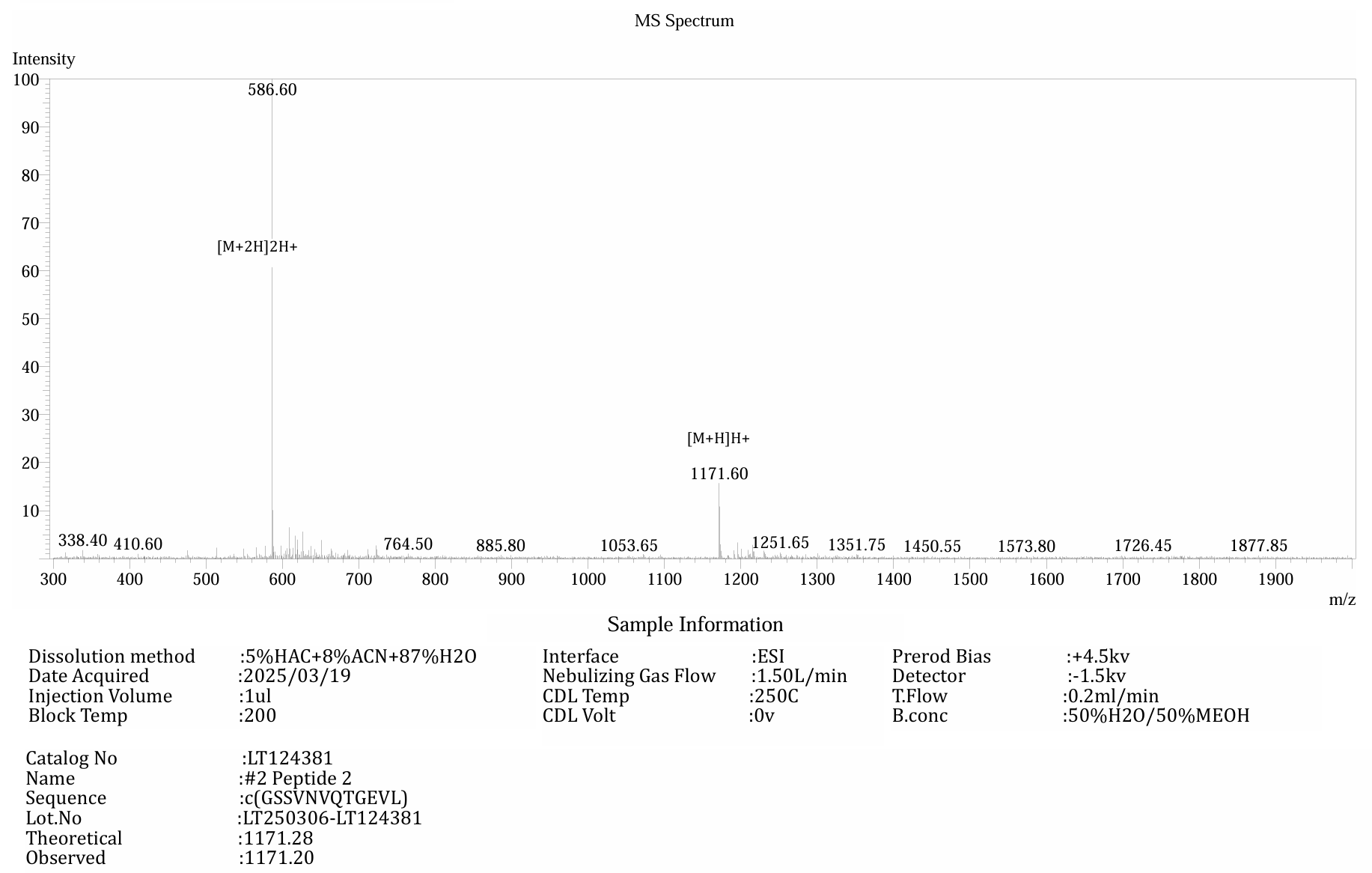

**Figure S3**. MS spectrum of **TP2**.

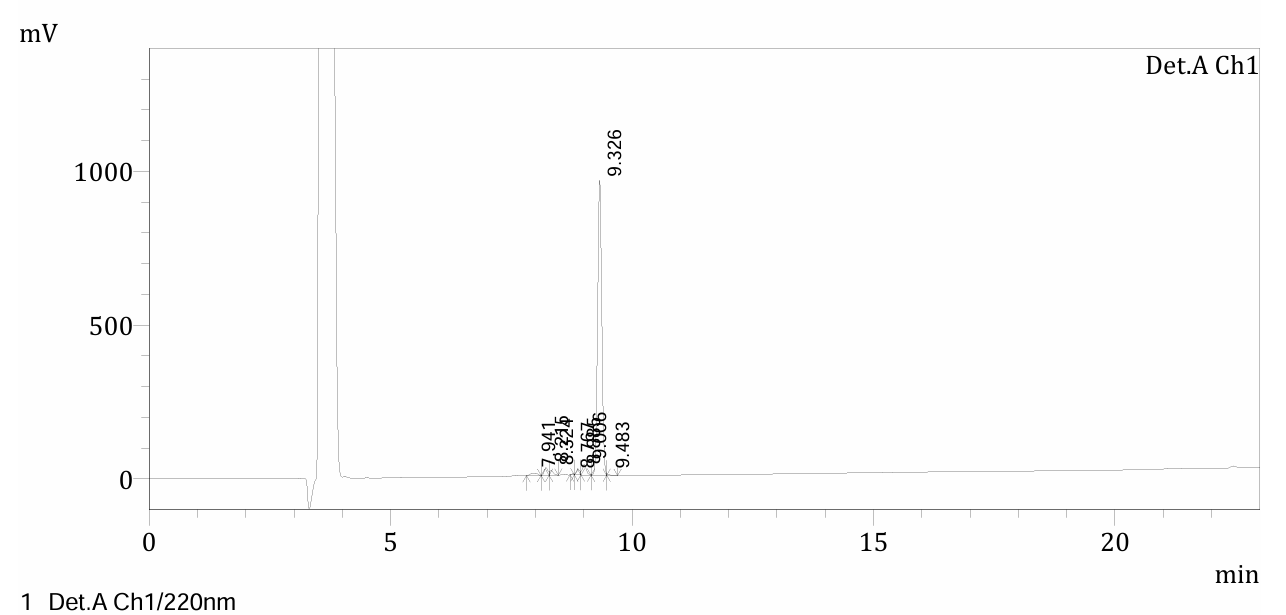

**Figure S4**. HPLC trace of **TP2**.

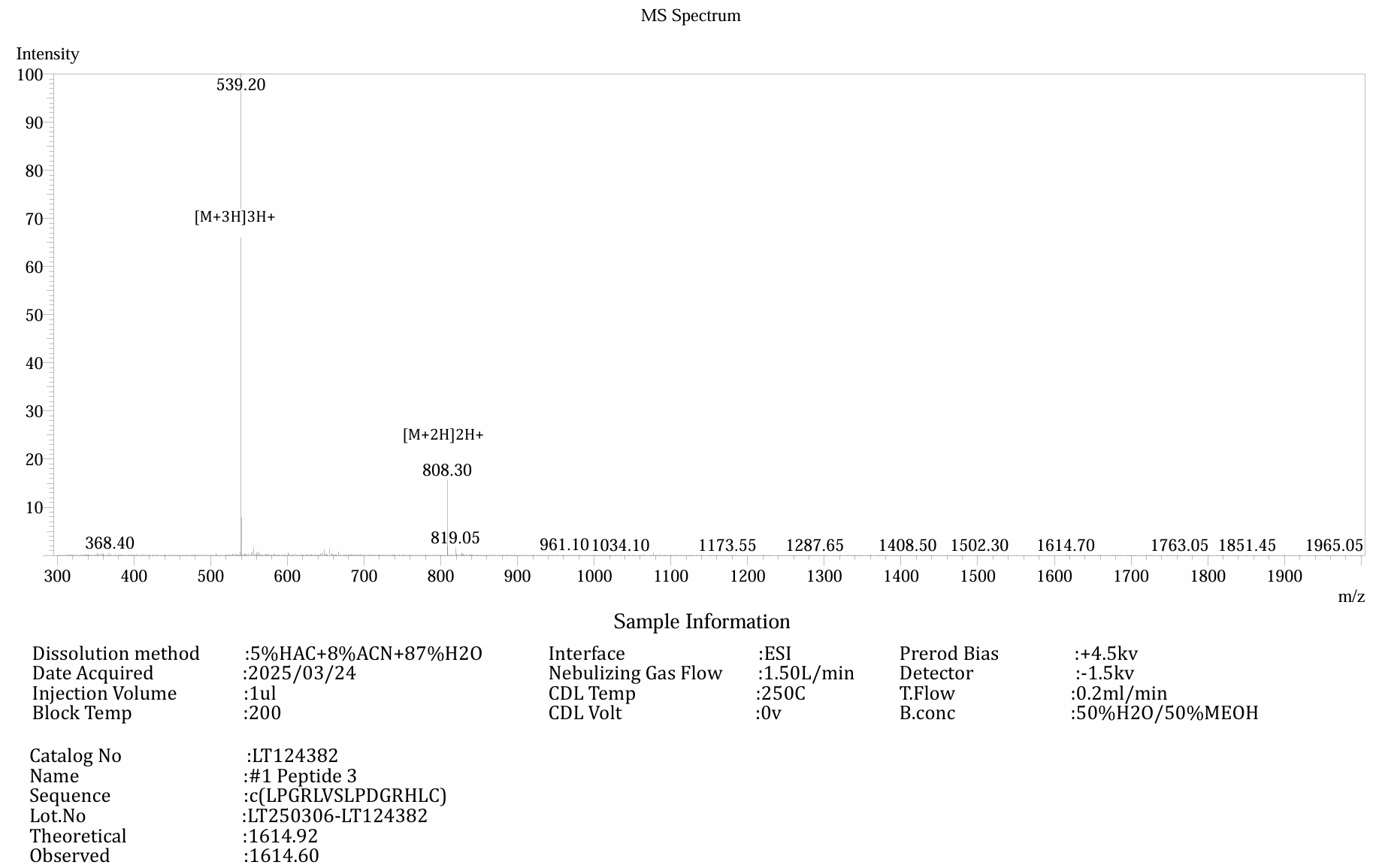

**Figure S5**. MS spectrum of **TP3**.

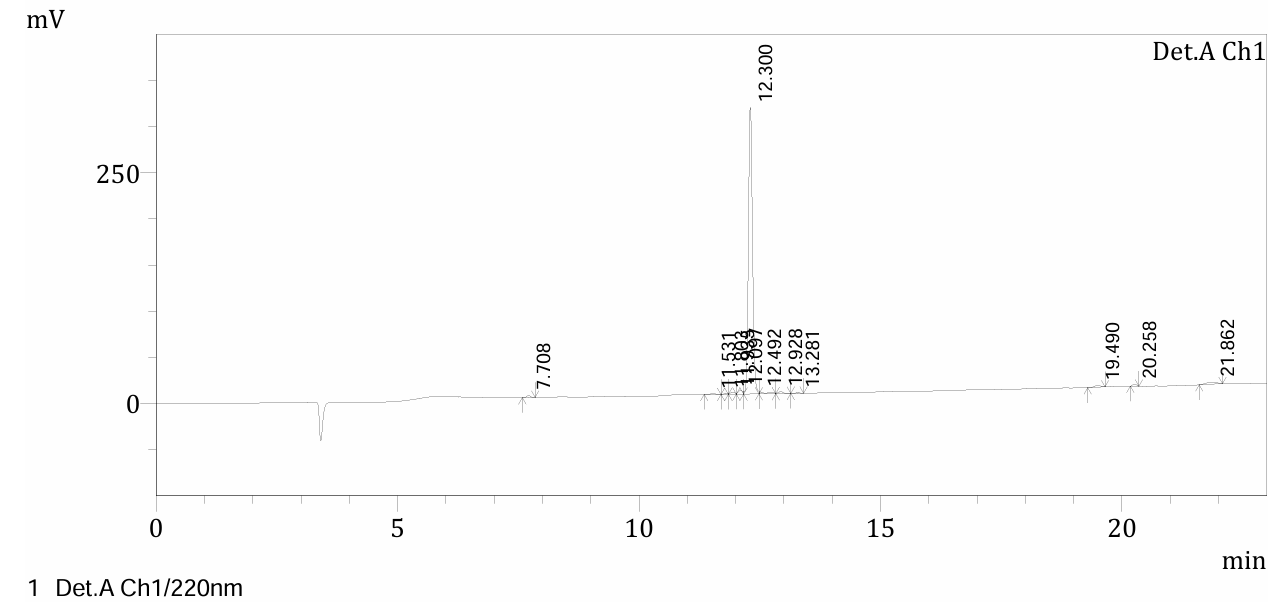

**Figure S6**. HPLC trace of **TP3**.

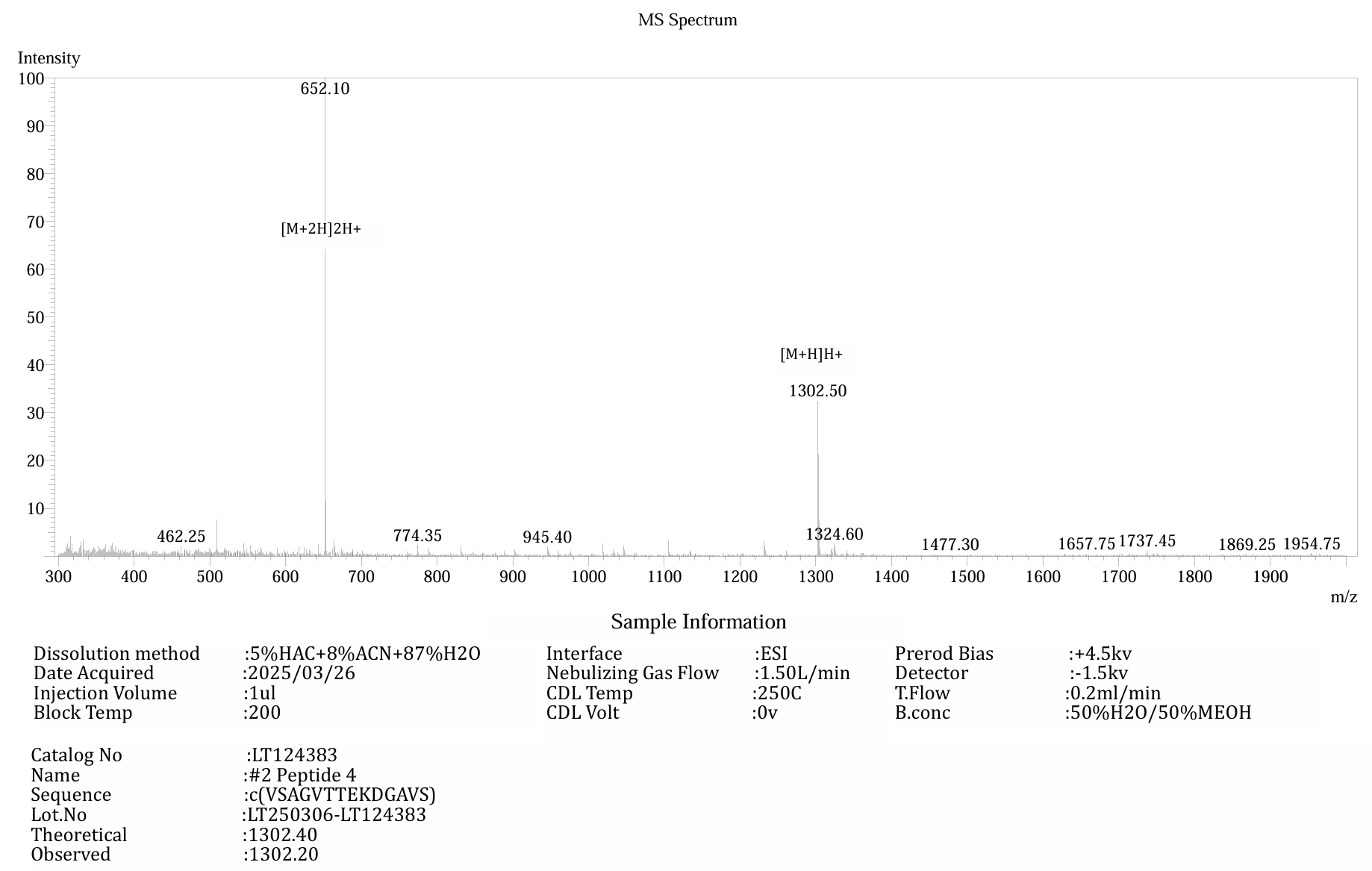

**Figure S7**. MS spectrum of **TP4**.

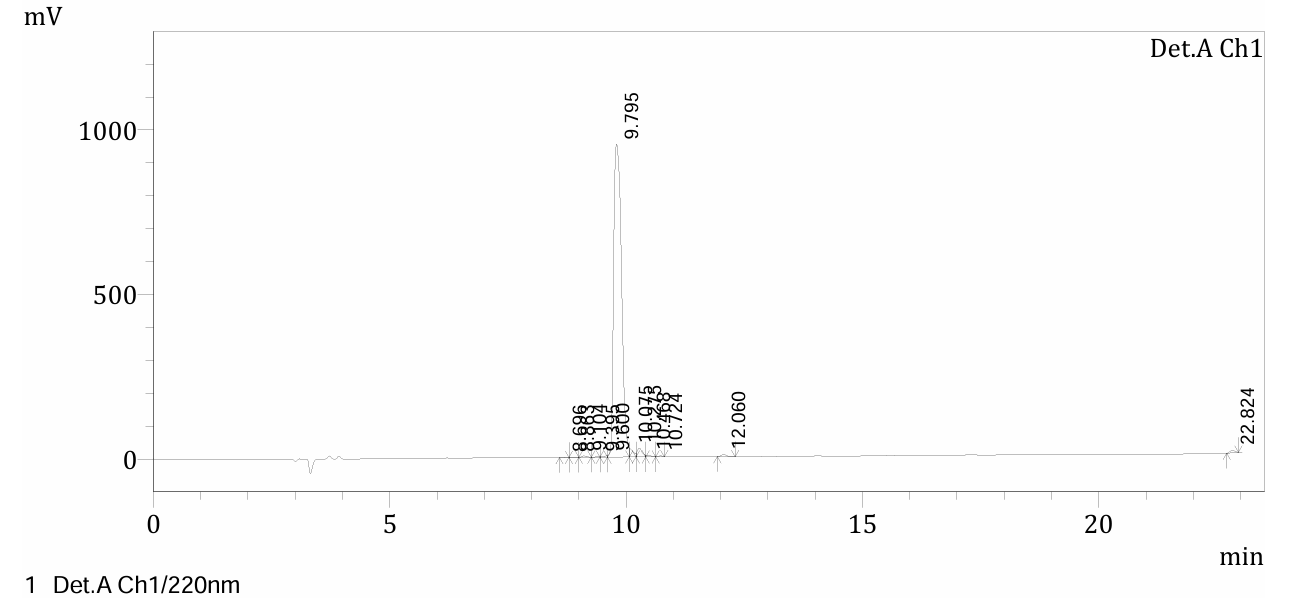

**Figure 8**. HPLC trace of **TP4**.
